# Development and characterization of Novel Small Molecule Inhibitors Targeting LAG-3 Protein

**DOI:** 10.1101/2025.08.29.673196

**Authors:** Oleksandr Yakovenko, Alireza Baradaran-Heravi, Steven J.M. Jones

## Abstract

Lymphocyte Activation Gene-3 (LAG-3) is a 503-amino acid transmembrane protein that modulates immune responses by negatively regulating the proliferation and activation of T cells - key effectors in adaptive immunity. The finely tuned expression of LAG-3, along with other immune checkpoints, prevents excessive immune activation and safeguards tissues from inflammation-induced damage. Importantly, the immune system also plays a critical role in tumor surveillance by recognizing and eliminating cells expressing neoantigens arising from somatic mutations. However, many cancers exploit immune checkpoint molecules like LAG-3 to dampen antitumor immunity. Elevated expression of LAG-3 within the tumor microenvironment contributes to immune evasion by suppressing cytotoxic T-cell activity. Consequently, inhibition of LAG-3 has emerged as a promising strategy for restoring immune function and enhancing anticancer immunity. This report presents the rational design and development of small-molecule inhibitors targeting LAG-3 through a novel semi-allosteric mechanism - *a priori* superior to classic (antibody) binding inhibitory - representing a next-generation therapeutic approach with potential applications in oncology and immune-related disorders.

## Introduction

The immune checkpoint receptor lymphocyte activation gene-3 (LAG-3/CD223; UniProtKB: P18627) was first identified in 1990 as a structural homolog of CD4 (1). It is expressed across diverse cellular populations, including both lymphocytic and non-lymphocytic lineages, where it functions as a negative regulator of antigen-specific T-cell activation and modulates their proliferation (2). LAG-3 exerts its inhibitory effects primarily through interactions with two major ligands: major histocompatibility complex class II (MHC II) (3) and fibrinogen-like protein 1 (FGL1) (4). Under physiological conditions, CD4 binds MHC II in concert with the T-cell receptor (TCR), facilitating T-cell activation via recruitment of Lck kinases, which phosphorylate components of the TCR complex (5). In contrast, LAG-3 not only competes with CD4 for MHC II binding, but also negatively regulates T cells by inhibiting CD3-mediated calcium flux and activation of the nuclear factor of activated T cells (NFAT) (6,7). Due to its immunosuppressive function, LAG-3 plays a critical role in the downregulation of immune responses in tumors and chronic viral infections. Notably, prior studies have demonstrated that CD4-dependent T-cell functionality can be restored using LAG-3-blocking immunotherapeutics (6,8,9). Encouraged by this therapeutic potential, over 20 LAG-3-targeting therapeutics are currently in development or undergoing various stages of FDA clinical trials.

Structurally, LAG-3 (CD223) is a transmembrane glycoprotein homologous to CD4, comprising four extracellular immunoglobulin-like domains (D1–D4) and a cytoplasmic tail involved in intracellular signaling (1). On the T-cell surface, LAG-3 forms dimers through interactions involving its D2 domain and transmembrane region associated with D4 (10,11). The N-terminal D1 domain serves as the primary ligand-binding interface, presenting two functionally distinct loops: the “extra loop” spanning amino acids A52–R76, which mediates high-affinity interaction with MHC class II molecules (10), and a second loop from G85–P93, which facilitates binding to fibrinogen-like protein 1 (FGL1). Intracellularly, LAG-3 transmits inhibitory signals via a membrane-proximal FXXL motif, the highly conserved ‘KIEELE’ sequence, and glutamic acid-proline (EP) repeats at the C-terminus (8-10). These motifs collectively suppress CD4-dependent T-cell activation by interfering with calcium flux and downstream transcriptional activity, including inhibition of nuclear factor of activated T cells (NFAT) signaling.

LAG-3 is widely recognized as the third clinically significant immune checkpoint, acting synergistically with programmed cell death protein 1 (PD-1) and cytotoxic T-lymphocyte antigen 4 (CTLA-4) (12-15). The US Food and Drug Administration (FDA) recently approved the anti-LAG-3 antibody relatlimab in combination with the anti–PD-1 antibody nivolumab for unresectable or metastatic melanoma (16), a regimen marketed under the name Opdualag (17). Relatlimab engages the D1 “extra loop” of LAG-3 by binding residues A59–W70, thereby blocking receptor–ligand interactions (18). Despite the impact of antibody-based checkpoint blockade, monoclonal therapies remain limited by parenteral administration, high manufacturing costs, and immune-related adverse events. Small-molecule inhibitors—offering oral bioavailability, tunable pharmacokinetics, and reduced immunogenicity—represent an attractive alternative; however, early-stage LAG-3 targeted compounds have not yet achieved adequate potency for clinical application (19). In this study, we describe the rational design and characterization of novel small-molecule semi-allosteric inhibitors of LAG-3 and demonstrate their competitive efficacy relative to relatlimab in a cell-based immunotherapy assay.

## Results

### Identification of the Small Molecule Inhibitor

#### Identification of a proper binding site on LAG-3 surface

To identify suitable small-molecule binding sites on LAG-3, we utilized the recently reported atomic structure of the extracellular region of human LAG-3 (PDB ID: 7TZG) (10). LAG-3’s extracellular segment comprises four immunoglobulin-like domains (D1–D4), with the D1 domain mediating high-affinity interactions with MHC class II ligands and the adjacent D2 domain contributing via spatial proximity and conformational support (20,21). Accordingly, our *in-silico* modeling efforts focused on a truncated construct containing only D1 and D2 domains. This approach substantially reduced the size of the periodic simulation box for molecular dynamics (MD) simulations while preserving the complete structural context of the LAG-3–MHC II interface.

To refine the experimental D1–D2 model of LAG-3, we employed a locally retrained version of the RoseTTAFold deep-learning framework (22) that incorporated the newest experimental LAG-3 coordinates into its training set. The initial atomic model produced by this network was then subjected to a 10 μs all-atom molecular dynamics (MD) simulation in explicit solvent to refine side chain packing and loop conformations. The fully equilibrated coordinates of the D1–D2 fragment are provided in Supplementary Information.

No small-molecule inhibitors of LAG-3 had been reported by that time, although cyclic nona-peptides C17 and C25 with approximately 1 µM potency were recently identified (23). To elucidate their binding modes, we performed extensive MD simulations using our refined D1–D2 LAG-3 model. For each peptide, we generated numerous random initial placements on the D1 surface and carried out 1 µs of explicit-solvent MD sampling for each. Peptide configurations that remained stably associated were then subjected to center-of-mass (COM) pulling to estimate their binding free-energy profiles. This protocol consistently revealed a high-affinity binding hotspot around residues R76 and Y77 – the sites previously shown experimentally to be critical for LAG-3 function (9). The conformation of the C17 peptide with the highest computed potential of mean force is detailed in Supplementary Information. This structure of LAG-3 bound to C17 complex subsequently served as the receptor conformation for our high-throughput virtual screening campaign.

#### High throughput virtual screening and hit identification

A diverse collection of ca. 70,000 compounds was obtained from ChemBridge Corporation, USA and screened virtually for potent compounds using the AutoDock Vina molecular docking software (24). Although Vina’s scoring function offers only an approximate ranking of binding affinities, it effectively enriches for true positives; therefore, we manually inspected the top 5 000 docked poses to select complexes with the most favorable orientations and scores. Each selected complex was subjected to a 1 μs explicit-solvent MD trajectory, and the resulting final coordinates were used for potential-of-mean-force (PMF) estimation via COM pulling. To broaden our hit set, we performed substructure searches for analogues of the highest-PMF compounds in the ChemBridge catalogue and evaluated these candidates with the same MD protocol, yielding a total of 320 unique protein–ligand complexes. From these, we ordered 32 compounds (10 %) based on PMF ranking and structural diversity to avoid redundancy. The inhibitory effects of the synthesized compounds on the LAG-3/MHCII interaction were assessed using the Homogeneous Time-Resolved Fluorescence (HTRF) LAG-3/MHCII binding assay (Revvity Inc., USA). Experimental screening of the purchased set identified LG-14, LG-16, and LG-17 (Figure 1) as LAG-3 inhibitors with *in vitro* potencies below 100 μM. Subsequent MD refinement of the LAG-3 bound to LG-17 complex (Figure 1) confirmed that LG-17 occupies the peptide-binding pocket around residues R76 and Y77, recapitulating the key interactions observed for the cyclic peptide inhibitors.

**Figure 1.**
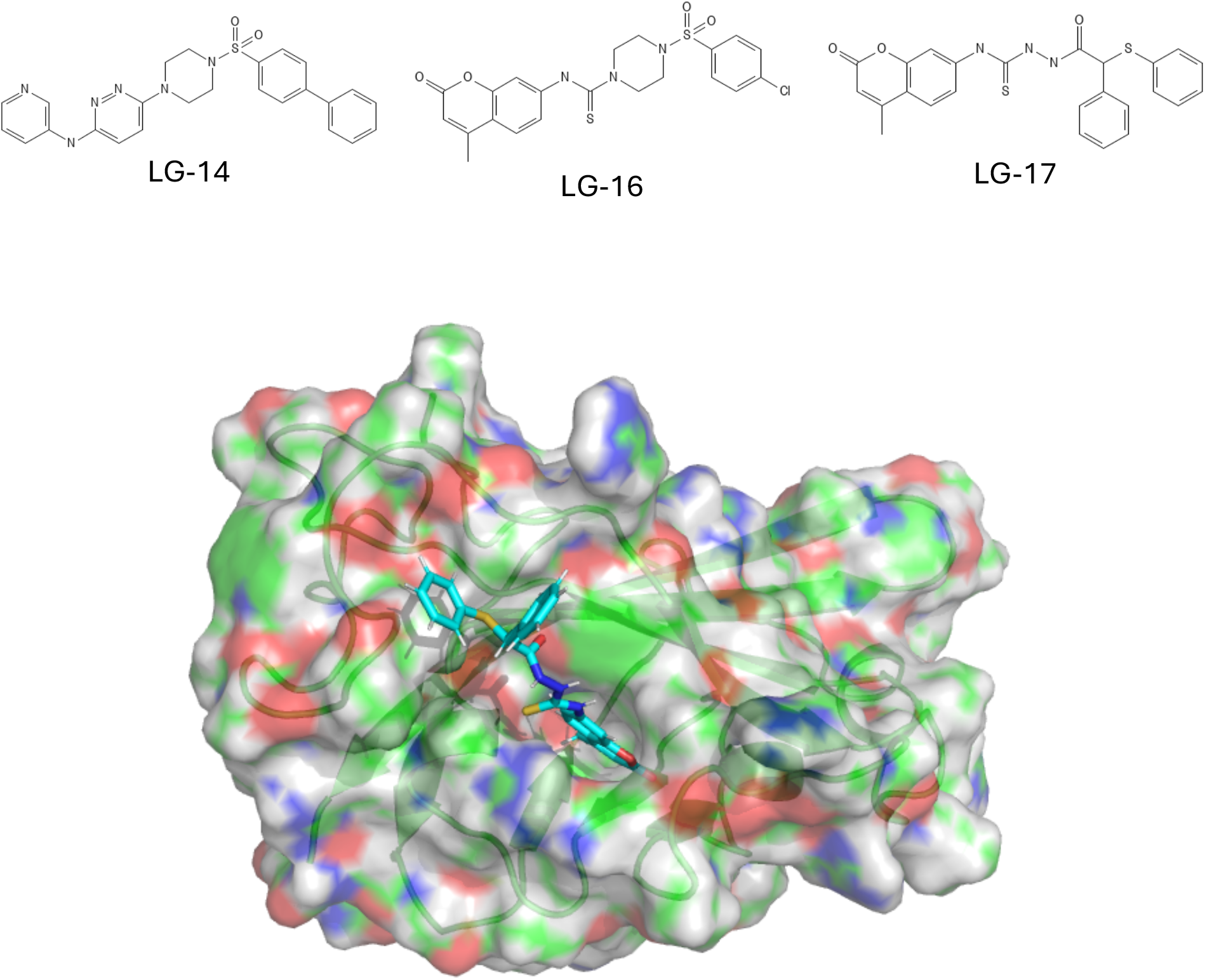
Structures of initial hits LG-14, LG-16 and LG-17 and the model of LAG-3 bound to LG-17 compound.

### Hit optimization

#### Scaffold design

Further optimization of LG-14, LG-16 and LG-17 hits were started with careful *in silico* design of the pharmacophore scaffold. We interpreted the results of experimental screening above as (4-methyl-) coumarin fragment to be the atomic subgraph which possess the promising activity of the tested inhibitors. As such, all our further derivatives were 4-methyl-coumarin derivatives. However, the coumarin fragment allows relatively small number of combinatorial chemistry opportunities, which are not always compatible with the structural models of LG-16 and LG-17 complexes with LAG-3 protein. For example, a series of virtual derivatives with substitution of the methyl in the fourth position of coumarin failed to show any improvements in MD PMF estimations.

Instead of further substituting chemically challenging coumarin fragment we appended, via a peptide bond, a phenyl ring with two chemically accessible branching spots in the second and fourth positions; such chemical architecture we refer as the core building block CBB0. The second position was designed as OH (referred to as OH-tail) and the fourth as NH2 (referred to as NH-tail). Such design allows flexible, controllable and relatively simple chemical modifications of each tail with desired building blocks. The design was specifically engineered to benefit from the parallel chemistry services provided by Enamine Inc., USA.

#### Round 1 of synthesis (Initial design of OH tail and in-wide screening of NH tail substitutions)

MD pulling simulations greatly supported the CBB0 design with typical PMF rising from ca. -30kJmol^-1^ of LG-14, LG-16, LG-17 hits to ca. -40kJmol^-1^ of virtual CBB0 derivatives. Because MD is a stochastic method, we did carry out a systematic replication of PMF estimations for the most potent virtual probes. The MD replication statistics showed that NH-tail substitutions have significantly higher PMF STD than OH-tail ones due to the “extra loop” of the D1 domain. The “extra loop” is an unstructured 30-residue insertion (A52–R76) which is unique to human LAG-3 within the immunoglobulin superfamily and contributes conformational flexibility at the ligand-binding interface (21). Indeed, structural and biochemical studies have demonstrated that this loop mediates interactions with MHC II molecules (25,26) constitutes the epitope targeted by the FDA-approved anti-LAG-3 antibody relatlimab (18).

In the MD-refined CBB0–LAG-3 complex, the appended NH-tail projects directly toward the N-terminal “extra loop” of the D1 domain, enabling transient contacts with loop residues. However, the intrinsic flexibility of this 30 amino-acid insertion produced a wide ensemble of conformations during MD trajectories, leading to large standard deviations in PMF estimates for NH-tail modifications. By contrast, the OH-tail consistently oriented toward a structurally stable β-sheet region in D2, yielding highly reproducible PMF free energies across replicate COM pulling simulations. Consequently, we elected to optimize the OH-tail *in silico* using MD-guided PMF analysis while reserving NH-tail diversification for empirical exploration via the Enamine Inc. parallel chemistry platform. A prototype benzamide substituent at the NH-tail established a virtual scaffold onto which over 100 distinct OH-tail analogues were manually appended and evaluated with identical MD-PMF protocols. Among these, derivatives bearing a benzoic acid motif achieved the most favorable binding free energies (ca. –50 kJ·mol−^1^). Synthesis and *in vitro* testing of compound LG-125, incorporating the benzamide NH-tail and benzoic acid OH-tail (Table 1), validated our predictions by delivering a 21 µM inhibitory potency. LG-125 therefore became the lead scaffold for subsequent NH-tail focused-lead optimization campaigns.

**Table 1.**
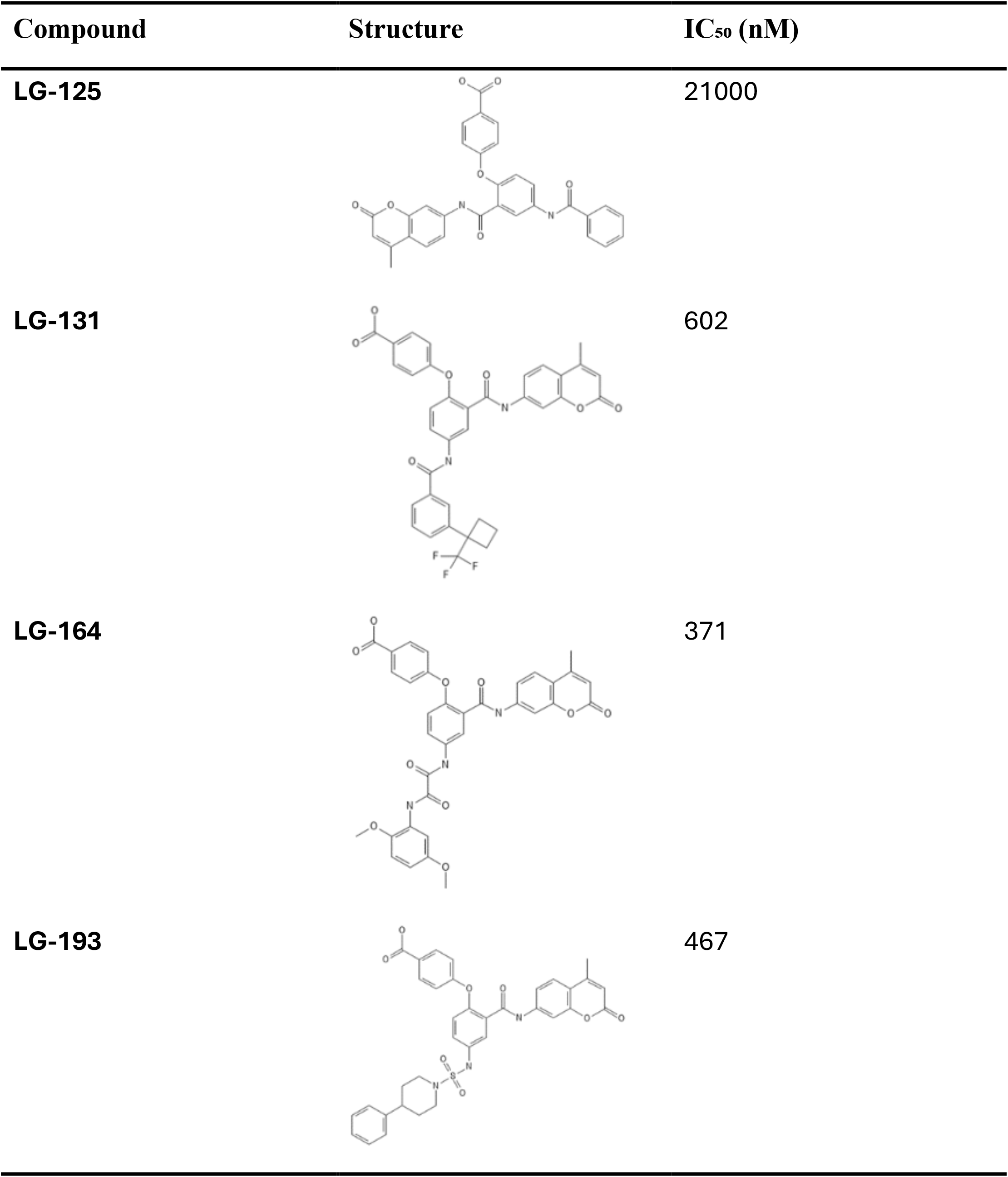
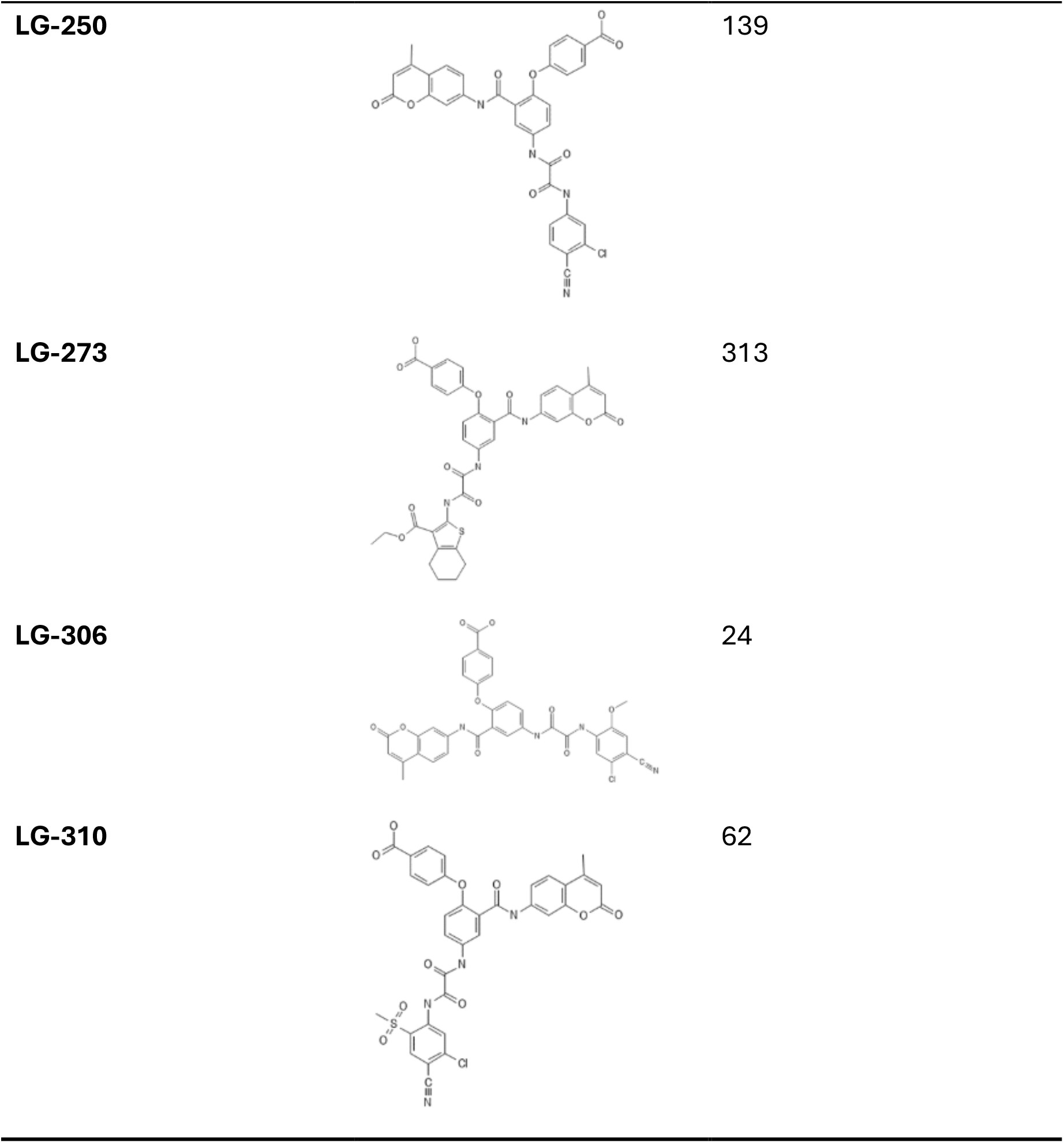
Structures and IC_50_ values of selected LAG-3 inhibitors.

| Compound | Structure | IC<sub>50</sub> (nM) |
| --- | --- | --- |
| LG-125 |  | 21000 |
| LG-131 |  | 602 |
| LG-164 |  | 371 |
| LG-193 |  | 467 |
| LG-250 |  | 139 |
| LG-273 |  | 313 |
| LG-306 |  | 24 |
| LG-310 |  | 62 |

In our first parallel synthesis campaign, we mined Enamine Inc’s building-block library to assemble 100 CBB0 derivatives that varied systematically in NH-tail linker length and conformational rigidity – the approach was driven by the challenges in reliably predicting NH-tail binding energetics with MD. Enamine Inc. chemists successfully synthesized 92 of these designs, and biochemical screening of the resulting library revealed three standout inhibitors: LG-131, LG-164, and LG-193 which each achieved sub-micromolar IC_50_ values (Table 1).

#### Round 2 of synthesis (Direct and semi-allosteric compounds, in-depth search of both tails)

In the second round of synthesis, we leveraged the diverse binding geometries observed for LG-131, LG-164, and LG-193 to guide a multi-directional optimization strategy. Structural analysis revealed three distinct semi-allosteric binding modes: LG-131, with its short, rigid amide linker, docks at the base of the “extra loop”; LG-164’s extended, stiff linker reaches directly into the loop region; and LG-193’s longer, more flexible linker engages an alternate surface on the D1 domain.

At this stage, we had formalized a predicted semi-allosteric inhibition model in which the benzoic acid–substituted coumarin OH-tail anchors competitively within LAG-3’s D1 binding cleft, while the NH-tail projects toward and allosterically modulates the dynamic “extra loop”. Our initial *in silico* evaluations, however, underestimated the contribution of LG-164’s extended linker and consequently prioritized LG-131 as the lead scaffold. We attribute this misassignment in part to the challenges of sampling the highly flexible “extra loop” by all-atom MD in practically reachable time horizon under conditions of throughput screening. As a result, we constructed a virtual library of 100 CBB0 derivatives drawn from Enamine Inc’s building-block collection and selected 100 compounds for synthesis, dividing them into two series: CBB1 analogues (inspired primarily by LG-131 but with inclusion of some close LG-164 structures) and CBB2 analogues (modeled on LG-193). During synthesis, the CBB1 series proved significantly more tractable from chemical perspective, yielding higher success rates and prompting a strategic shift toward amide-linked scaffolds for downstream optimization.

Biochemical profiling of 82 newly synthesized compounds from our second parallel chemistry identified two outstanding candidates – LG-250 and LG-273 both of which belong to LG-164 structures series and exhibit increased inhibitory potency in *in vitro* assays (Table 1).

#### Round 3 of synthesis (Iterative optimization and synthesis)

In the final synthetic round, we focused exclusively on optimizing our semi-allosteric inhibitors. The superior potency of LG-250 enabled us to refine our allosteric model of extra-loop folding (Figure 2). By incorporating simulations of the inactive loop conformation, we could accurately predict each compound’s dual contributions – competitive anchoring and allosteric modulation – to overall inhibitory potency. Accordingly, we conducted several iterative cycles of intensive molecular-dynamics simulation to tailor analogues of LG-250 and LG-273 for their ability to induce and stabilize the inactive extra-loop conformation of LAG-3. Top MD-ranked candidates were synthesized by Enamine Inc. Although a subset of the proposed analogues proved synthetically too challenging, 11 new compounds were delivered. Among these, LG-306 and LG-310 emerged as deep-nanomolar inhibitors in our *in vitro* assay (Table 1).

**Figure 2.**
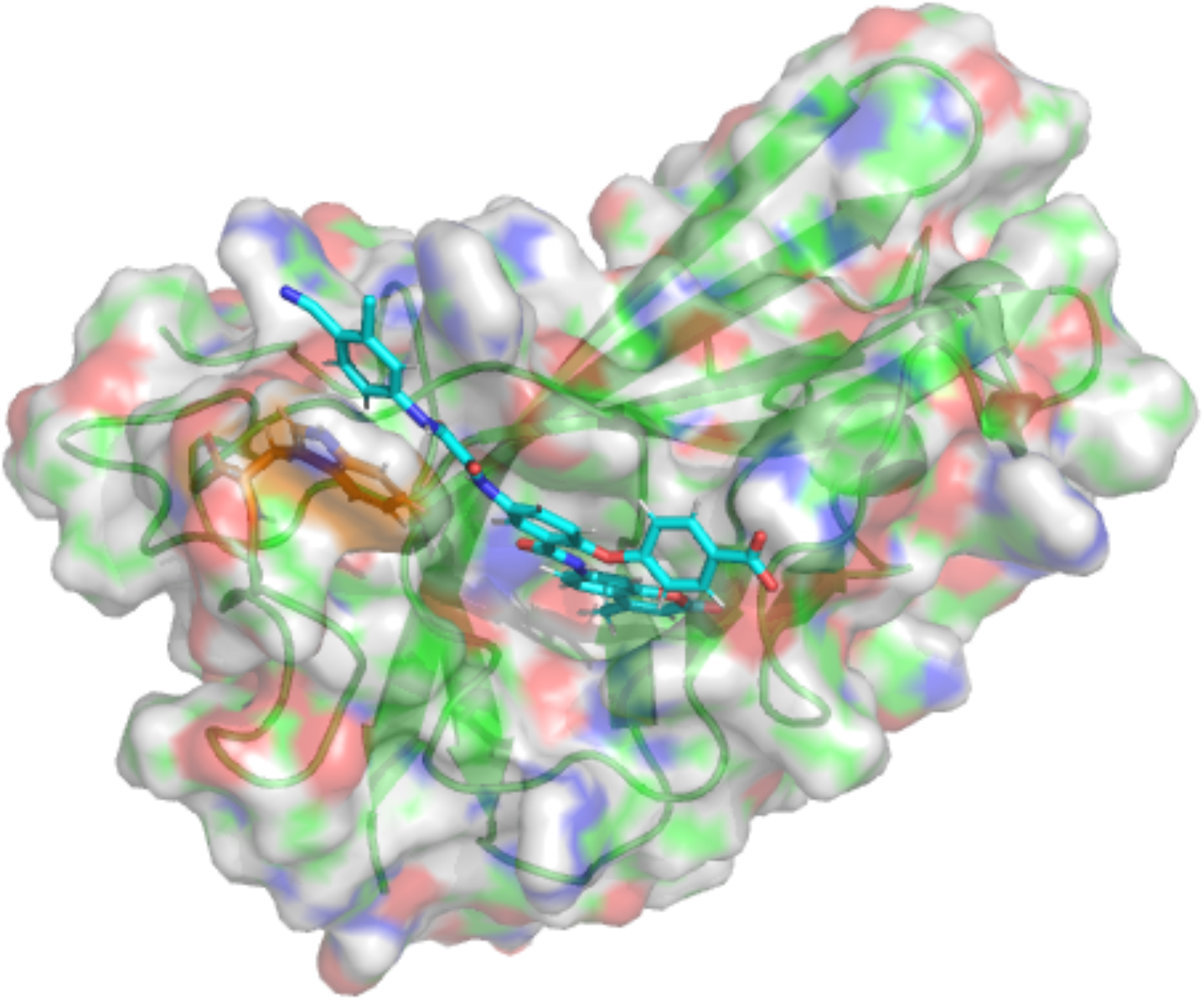
Model of D1 domain of LAG-3 in complex with the semi-allosteric inhibitor LG-250. The orange sticks show capture of Trp70 in the middle of the “extra loop”. The D2 domain is hidden for clarity.

### Cellular assay demonstrates advantages of semi-allosteric compounds

Compounds that demonstrated the strongest potency in *in vitro* experiments were evaluated in a cell-based assay designed to mimic the immune response, providing an integrated readout of complex, multifactorial interactions. This assay consists of an MHCII-positive cell line and Jurkat T cells engineered to express human LAG-3 along with a luciferase reporter under the control of T-cell activation pathway-dependent response elements. When co-cultured, the MHCII-positive cells present a T-cell receptor (TCR)-activating antigen to Jurkat T cells, triggering TCR signaling and promoter-driven luminescence. However, LAG-3 engagement with MHCII suppresses TCR signaling, leading to a reduction in luminescence.

Compounds or antibodies that disrupt LAG-3/MHCII binding restore TCR pathway activation, resulting in a dose-dependent increase in luminescence. Some candidates, particularly LG-164, LG-306 and LG-310 induced measurable T-cell activation at concentrations above 1 µM (Figure 3). Interestingly, we observed that at saturating doses, at least one semi-allosteric compound, LG-310, reached and even outperformed the commercial anti-LAG-3 antibodies, Clone 17B4 and relatlimab, in T-cell activation. We attribute this superior maximal efficacy of our drug candidate to the semi-allosteric mechanism of its action.

**Figure 3.**
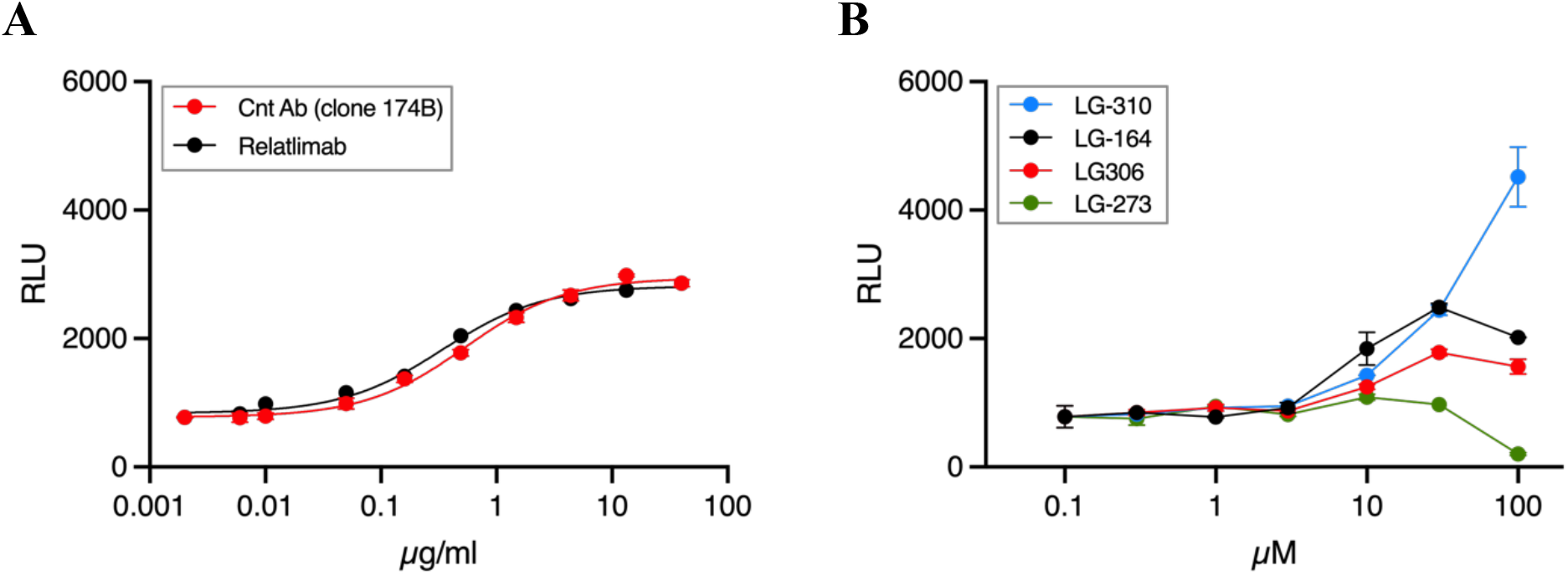
Activity in cellular assay for the control antibody clone 17B4 (Promega Inc) and FDA approved Relatlimab (Selleck Chemicals LLC) (**A**); the activities of discovered small organic compounds (**B**).

## Discussion

### Structural basis for specificity and potency

Structural determinants of specificity and potency became evident through extensive MD simulations starting from LG-16 and LG-17 bound to the LAG-3 D1–D2 fragment (Figure 1). In these models, the (4-methyl-)coumarin core consistently docks into a hydrophobic pocket formed predominantly by Loop 2 and centered on Y77, where π– π stacking secures the ligand. This anchoring motif, preserved across our lead series, confers selectivity for LAG-3 over other immunoglobulin-superfamily receptors. The oxanilide linker, by virtue of its length and rigidity, projects the NH-tail farther toward the unique structural element “extra” loop of LAG-3, specifically positioning it to interact with W70 and stabilize a non–binding conformation of the loop. The W70 residue is by far the most hydrophobic component of the unstructured “extra” loop and its stabilization can be the key step in promoting an artificially folded state for this structural fragment. As a result, the inhibitors which stabilize W70 exert a dual action: orthosteric competition at the Loop 2 pocket disrupts FGL1 engagement, while allosteric locking of “extra” loop prevents strong MHC II binding.

Although primary focus of LAG-3 inhibitors design in our study was at the predicted semi-allosteric mechanism of action, we also speculate that well-tailored OH-substituents of benzoic acid would further enhance Loop 2 interactions, strengthen the potency of compounds and even modulate LAG-3 interactions with FGL1. Collectively, these insights have guided the development of a new family of multi-modal, highly selective LAG-3 inhibitors.

### Semi-allosteric mechanism of action

Theoretical molecular modelling of LG-250 bound to LAG-3 (Figure 2) reveals a dual engagement that underlies its semi-allosteric mode of inhibition. In the refined LAG-3 bound to LG-250 complex, the ortho-chlorocyanobenzene moiety of the inhibitor docks against the indole side chain of W70, a residue centrally located within the “extra loop” (A52–R76) known to mediate MHC II binding. This contact induces a conformational shift in the extra loop, displacing it from its canonical trajectory toward the MHC II interface. Although atomistic simulations of the LAG-3 bound to MHC II complex were beyond this study’s scope, we hypothesize that the remodeled loop cannot adopt the geometry required for productive MHC II engagement. Hence, we predict LG-250 combines direct competitive anchoring in the D1 binding cleft with stabilization of an inactive “extra” loop conformation. We define this dual mechanism as semi-allosteric inhibition, where allosteric loop locking augments, but does not replace the inhibitor’s orthosteric blockade of LAG-3.

The semi-allosteric model is borne out by our cell-based assays: compounds LG-164, LG-250, LG-273, LG-306, and LG-310 each restored T-cell activation in a physiological context, whereas LG-131 and LG-193, despite its sub-micromolar IC_50_ values in biochemical screens, failed to inhibit LAG-3 signaling in cell tests. These findings imply that the LG-250 series operates via a dual mechanism: the benzoic-acid–coumarin core anchors competitively in the D1 cleft, while the NH-tail allosterically locks the “extra loop” into an inactive conformation. By contrast, LG-131 and LG-193 rely solely on orthosteric blockade of the binding surface, a strategy sufficient to displace small peptide ligands *in vitro* but it is insufficient to suppress it in the more complex cellular environment.

Another line of evidence supporting the proposed semi-allosteric mechanism is the observation of maximal potency at the saturating concentrations (Figure 3). Notably, LG-310, one of the most potent compounds in the series, significantly exceeds the efficacy achieved by antibody-based blockade. This enhanced activity suggests that LG-310’s mechanism extends beyond a simple orthosteric inhibition. We attribute this additional functionality to compound-induced conformational remodeling of the “extra loop,” which likely stabilizes LAG-3 in an inactive state. Importantly, this alternative loop folding may exhibit hysteresis, keeping LAG-3 in a non-functional conformation even after the compound dissociates from the binding pocket.

### Advantages compared with existing therapies

Compared with FDA approved antibody-based checkpoint blockade, our semi-allosteric LAG-3 inhibitors offer several therapeutic advantages. First, compound LG-310 achieves nanomolar potency and, in cell-based assay, exceeds the functional inhibition of relatlimab at saturation. The small-molecule nature confers oral bioavailability, tunable pharmacokinetics, and tissue penetration—attributes that antibodies lack. Moreover, the dual mechanism of action, combining orthosteric anchoring with allosteric “extra” loop locking, provides a more durable receptor inactivation without the immunogenicity or infusion-related adverse events associated with monoclonal antibodies. Taken together, this proposed semi-allosteric scaffold not only matches the efficacy of current LAG-3 antibodies but also promises a more flexible and patient-friendly therapeutic profile. It will be important to further optimize the compounds and test them in humanized animal models to determine whether therapeutically relevant LAG-3 inhibition and tumor suppression can be achieved with a good therapeutic window *in vivo*.

In conclusion, the discovery and optimization of semi-allosteric small-molecule LAG-3 inhibitors provide an advance toward potential orally bioavailable checkpoint modulators with antibody-like efficacy. By uniting competitive anchoring in the extracellular binding cleft with loop-locking allosteric regulation, these compounds achieve nanomolar potency and inhibition of cellular activity – surpassing commercial antibodies at saturating doses – while offering the pharmacokinetic flexibility and reduced immunogenicity inherent to small molecules. Importantly, this proposed dual-mechanism design framework is readily adaptable to other immune checkpoints, such as CTLA-4, laying the groundwork for combinable, multi-targeted immunotherapies that sidestep the manufacturing, delivery, and safety challenges of monoclonal antibodies.

## Methods

### In-silico Screening and Compound Selection

Autodock Vina free molecular docking software (24) was used for high-throughput virtual screening of large collections of small organic compounds. Under its default settings Vina is capable of processing a library of >100K candidates in less than one week with one multicore AMD Threadripper workstation. Large-scale, GPU-enabled simulations of molecular dynamics (MD) were employed for more precise evaluation of the binding strength between the LAG-3 protein and a selected small organic compound candidate. All MD simulations were carried out using GROMACS-2020 package (27,28) with AMBER99SB-ILDN fore field (29). The estimations of the potential of mean force (PMF) over the dissociation reaction coordinate were computed with umbrella sampling method. Analysis of the pulling trajectories was performed with the weighted histogram analysis method (WHAM) (30). Our MD screening protocol is composed of several stages as follows. A model of the protein bound to a candidate compound is placed into a 9×9×9 nm cubic periodic box of TIP3P water media. The system is equilibrated. Firstly, with a short constant volume warming to T=310K simulation followed by a short constant pressure relaxation at one atmosphere. Secondly, a long – 256 ns – NPT simulation is carried out to sample the adaptive conformational changes in the protein induced by the presence of a compound candidate. The long equilibration also assured the stability of candidate complexes under physiological temperature and pressure. For the stable complexes an artificial pulling force of 1000 KJmol^-1^nm^-1^ s applied along the dissociation reaction coordinate for 16 ns to generate a dissociation trajectory. The trajectory is then sliced into 12 umbrella windows at 0.0, 0.1, 0.2, 0.3, 0.4, 0.5, 0.6, 0.7, 0.8, 0.9, 1.0 and 1.2 nm of center of mass separation and each window is sampled for 32 ns long productive run with umbrella potential locked at the windows centers. The resulting trajectories are fitted with WHAM to reconstruct the reactions’ Potential of Mean Force (PMF), but the first 2 ns of them were systematically excluded from the analysis to allow for equilibration after the non-equilibrium dissociation pulling. The adequate parameters of small organic compounds for the simulations were produced by our in-house implementation (31-33).

## Chemical Synthesis

### Synthesis of tert-butyl 4-(4-amino-2-((4-methyl-2-oxo-2H-chromen-7-yl)carbamoyl)-phenoxy)-benzoate (CBB)

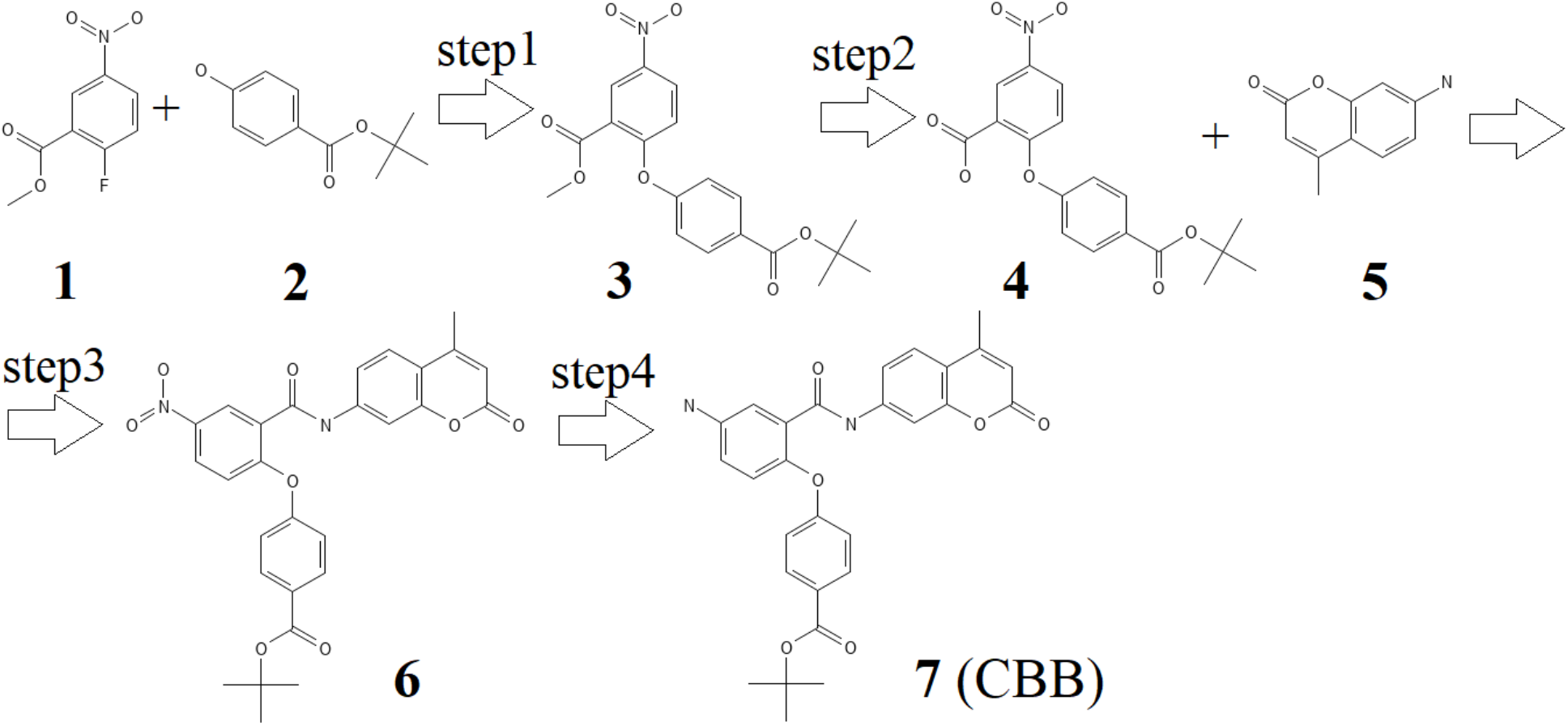

### Step1 of CBB synthesis

Synthesis of methyl 2-(4-(tert-butoxycarbonyl)phenoxy)-5-nitrobenzoate (3). A mixture of compound 1 (44.8 g, 0.225 mol), compound 2 (43.7 g, 0.225 mol), and K2CO3 (46.6 g, 0.337 mmol) in DMF was heated at 80°C for 16 h. The mixture was diluted with water, extracted with EtOAc (1 L × 3), the organic layer was washed water (200 mL × 2) and brine (100 mL), dried over N2SO4, filtered, and evaporated to give compound 3 (70.5 g, 0.189 mol, 84% yield).

### Step2 of CBB synthesis

Synthesis of 2-(4-(tert-butoxycarbonyl)phenoxy)-5-nitrobenzoic acid (4). A mixture of compound 3 (70.5 g, 0.189 mol) and LiOH monohydrate (9.5 g, 0.227 mol) in THF: water was stirred at room temperature for 16 h. Next, it was concentrated in vacuo, acidified to pH=4, and extracted with EtOAc (700 mL × 3), the organic layer was washed water (200 mL × 2) and brine, dried over Na2SO4, filtered, and evaporated to give compound 4 (60.3 g, 0.168 mol, 88.9% yield).

### Step3 of CBB synthesis

Synthesis of tert-butyl 4-(2-((4-methyl-2-oxo-2H-chromen-7-yl)carbamoyl)-4-nitrophenoxy)benzoate (6). To a solution of compound 4 (60.3 g, 0.168 mol) and compound 5 (24.5 g, 0.140 mol) in DMF was added DIPEA (97.44 mL, 0.559 mmol) and HATU (69.13 g, 0.182 mmol) at 0 °C, and the mixture was stirred overnight at rt. The mixture was diluted with water, and the precipitate was collected by filtration, washed with water three times and MTBE, dried to give compound 6 49.3 g (95.4 mmol, 68.2 %).

### Step4 of CBB synthesis

Synthesis of tert-butyl 4-(4-amino-2-((4-methyl-2-oxo-2H-chromen-7-yl)carbamoyl)phenoxy)benzoate (CBB). A solution of compound 6 (25 g, 48.4 mmol) and 4,4’-bipyridine (0.38 g, 2.42 mmol) in anhydrous DMF was cooled down to -10 °C. Then hypoboric acid (13 g, 145 mmol) was added in three portions over 30 min. The reaction mixture was stirred at ambient temperature for another 30 minutes and then poured into an ice-cold saturated K2CO3 aqueous solution. The mixture was diluted with water, and the precipitate was collected by filtration, washed with water three times, and MTBE, dried to give CBB (20 g, 41.1 mmol, 84.9%). ^1^H NMR (500 MHz, DMSO-*d*_6_) δ 10.59 (s, 1H), 7.76 (d, *J* = 8.9 Hz, 2H), 7.66 (d, *J* = 8.3 Hz, 2H), 7.53 (d, *J* = 8.7 Hz, 1H), 6.92 – 6.84 (m, 4H), 6.75 (dd, *J* = 8.6, 2.8 Hz, 1H), 6.24 (s, 1H), 5.37 (s, 2H), 2.36 (s, 3H), 1.47 (s, 9H).

### CBB Amidation and Deprotection

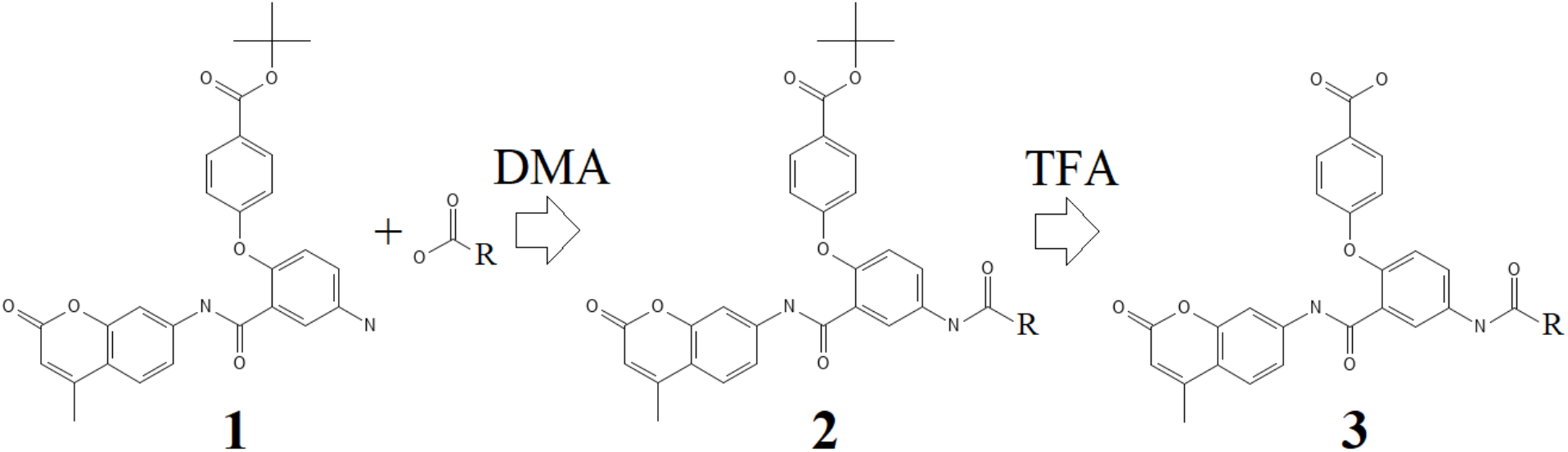

Reagent 1 – CBB – (1 eq.), Reagent B (1.3 eq.), HATU (1.5 eq.), and DIPEA (2.5 eq.) were mixed in dry DMA (appr. 0.7 ml per 100 mg of product). In case of using a salt of any of reagents, an additional amount of DIPEA was added to the reaction mixture to transfer the reagent to the base form. The mixture was sealed and stirred at 60 C for 16 hour(-s). Then the solvent was evaporated under reduce pressure and the cleavage cocktail (trifluoroacetic acid, triisopropylsilane, water (93:5:2; v/v) appr. 0.7 ml per 100 mg of product) was added to the mixture. The mixture was stirred at ambient temperature for 6 hour(-s). The solvent was evaporated under reduced pressure, and the residue was dissolved in DMSO (appr. 1 ml up to 100 mg of product). The solution was filtered, analyzed by LCMS, and transferred for HPLC purification.

The purification was performed using Agilent 1260 Infinity systems equipped with DAD and mass-detector. Waters Sunfire C18 OBD Prep Column, 100 A, 5 µm, 19 mm × 100 mm with SunFire C18 Prep Guard Cartridge, 100 A, 10 µm, 19 mm × 10 mm was used. Deionized Water (phase A) and HPLC-grade Methanol or Acetonitrile (phase B) were used as an eluent. In some cases, ammonia or TFA was used as an additive to improve the separation of the products. In these cases, free bases and TFA salts of the products were formed respectively.

### CBB Sulfonamidation (RSO_2_Cl) and Deprotection

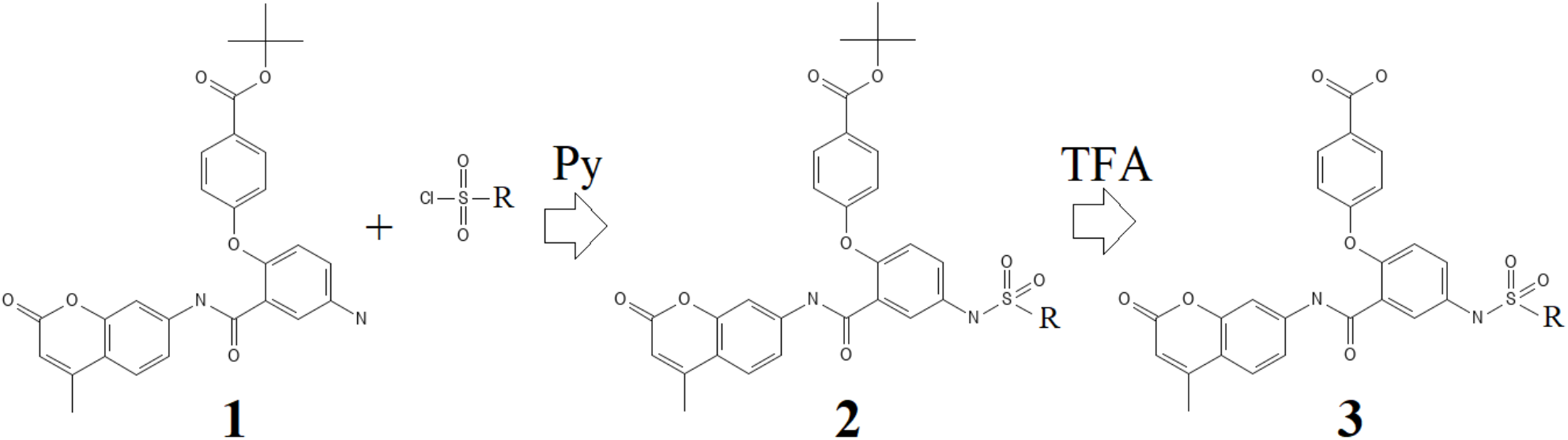

Reagent 1 – CBB – (1 eq.), Reagent B (1.2 eq.), DMAP (0.05 eq.), were mixed in dry Py (appr. 2 ml per 100 mg of product). The mixture was sealed and stirred at ambient temperature for 24 hour(-s). Then the solvent was evaporated under reduce pressure and the cleavage cocktail (trifluoroacetic acid, methylene chloride (15:100; v/v) appr. 2 ml per 100 mg of product) was added to the mixture. The mixture was stirred at ambient temperature for 72 hour(-s). The solvent was evaporated under reduced pressure, and the residue was dissolved in DMSO (appr. 1 ml up to 100 mg of product). The solution was filtered, analyzed by LCMS, and transferred for HPLC purification.

The purification was performed using Agilent 1260 Infinity systems equipped with DAD and mass-detector. Waters Sunfire C18 OBD Prep Column, 100 A, 5 µm, 19 mm × 100 mm with SunFire C18 Prep Guard Cartridge, 100 A, 10 µm, 19 mm × 10 mm was used. Deionized Water (phase A) and HPLC-grade Methanol or Acetonitrile (phase B) were used as an eluent. In some cases, ammonia or TFA was used as an additive to improve the separation of the products. In these cases, free bases and TFA salts of the products were formed respectively.

### Synthesis of 5-amino-2-hydroxy-N-(4-methyl-2-oxo-2H-chromen-7-yl)benzamide (CBB0)

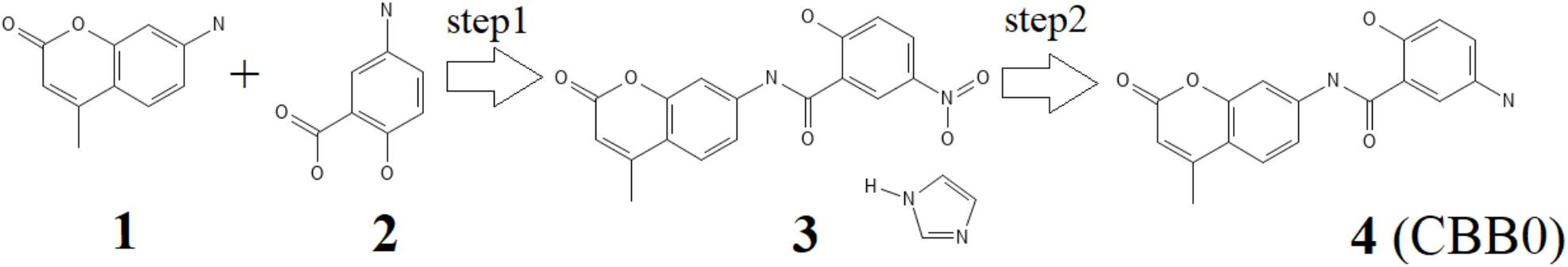

### Step1 of CBB0 synthesis

Synthesis of 1H-imidazol-3-ium 2-[(4-methyl-2-oxo-2H-chromen-7-yl)carbamoyl]-4-nitrobenzen-1-olate (3): 2-Hydroxy-5-nitrobenzoic acid (17.5 g, 95.62 mmol, 1 eq.) was dissolved in dry dioxane (600 ml). Then 1-(1H-imidazole-1-carbonyl)-1H-imidazole (16.27 g, 100.4 mmol, 1.05 eq.) was added and the resulting mixture was stirred at 50°C for 1 h. 7-Amino-4-methyl-2H-chromen-2-one (16.74 g, 95.62 mmol, 1 eq.) was dissolved in dioxane (600 ml) under heating. The resulting solution was added to the reaction mixture and the resulting mixture was stirred at 100°C for 12 h. Then the reaction mixture was cooled to ambient temperature and filtered off. The filter cake was then washed with MTBE (3 x 150 ml) and dried in vacuo to give 1H-imidazol-3-ium 2-[(4-methyl-2-oxo-2H-chromen-7-yl)carbamoyl]-4-nitrobenzen-1-olate (22.3 g, 95.0% purity, 51.88 mmol, 54.3% yield) as a yellow solid.

^1^H NMR (500 MHz, DMSO) δ 14.30 (s, 1H), 8.75 (s, 1H), 8.71 (s, 1H), 7.98 (s, 1H), 7.89 (dd, J = 9.6, 3.3 Hz, 1H), 7.71 (d, J = 8.6 Hz, 1H), 7.53 (s, 2H), 7.40 (d, J = 8.5 Hz, 1H), 6.43 (d, J = 9.5 Hz, 1H), 6.23 (s, 1H), 2.40 (s, 3H).

### Step2 of CBB0 synthesis

Synthesis of 5-amino-2-hydroxy-N-(4-methyl-2-oxo-2H-chromen-7-yl)benzamide (CBB0): 1H-imidazol-3-ium 2-[(4-methyl-2-oxo-2H-chromen-7-yl)carbamoyl]-4-nitrobenzen-1-olate (10.0 g, 24.5 mmol, 1 eq.) was suspended in dioxane/H_2_O mixture (1:1, 400 ml). Sodium hydrosulfite (42.61 g, 245.02 mmol, 10 eq.) was added and reaction mixture was stirred at 110°C for 72 h. After confirmation of completion of the reaction by LCMS the reaction mixture was cooled to ambient temperature and filtered off. The filter cake was washed with H_2_O (2 x 100 ml) and dioxane (2 x 100 ml). The solid residue was dried in vacuo to give 5-amino-2-hydroxy-N-(4-methyl-2-oxo-2H-chromen-7-yl)benzamide (6.0 g, 95.0% purity, 18.37 mmol, 75% yield) as a yellow solid.

^1^H NMR (500 MHz, DMSO) δ 10.76 (s, 1H), 10.02 (s, 3H), 7.96 (s, 1H), 7.88 – 7.74 (m, 2H), 7.67 (d, J = 8.7 Hz, 1H), 7.40 (d, J = 8.7 Hz, 1H), 7.17 (d, J = 8.6 Hz, 1H), 6.34 (s, 1H), 2.46 (s, 3H)

### The Synthesis of 2-hydroxy-N-(4-methyl-2-oxo-2H-chromen-7-yl)-5-(3-(1-(trifluoromethyl)cyclobutyl)-benzamido)benzamide (CBB1)

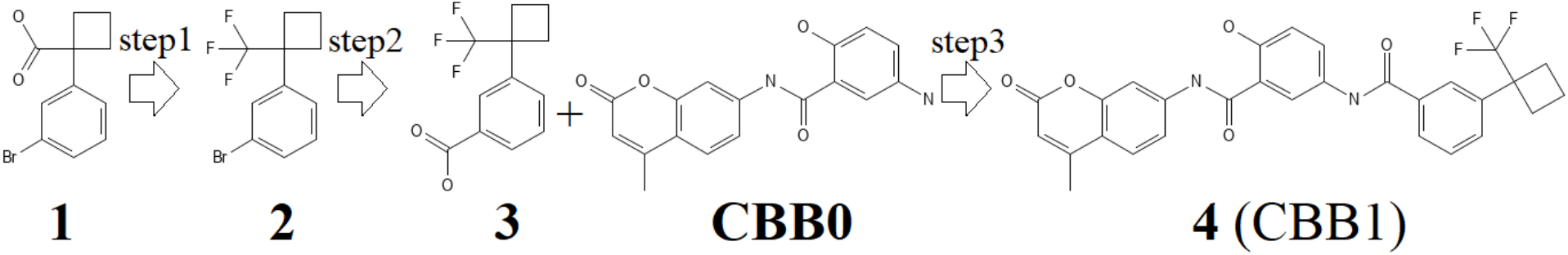

### Step 1 of CBB1 synthesis

The Synthesis of 1-bromo-3-[1-(trifluoromethyl)cyclobutyl]benzene (2): 1-(3-Bromophenyl)cyclobutane-1-carboxylic acid (5.0 g, 19.69 mmol) was placed in autoclave. The reaction vessel was cooled down by liquid nitrogen to -100°C and anhydrous hydrogen fluoride (5.91 g, 295.27 mmol, 15 eq.) was added. Then sulfur tetrafluoride (6.38 g, 59.05 mmol, 3 eq.) was condensed into the reaction vessel. Cooling bath was removed, and the mixture was allowed to warm up to a room temperature. It was then heated at 60°C in an oil bath for 72 h. The autoclave was allowed to cool down to room temperature, and the gaseous products were vented off into a trap with an aqueous solution of NaOH (1 M). The residue was poured onto ice (50 g), neutralized with a saturated aqueous solution of NaHCO_3_ to pH = 8. The solution was extracted with CH_2_Cl_2_ (3 × 50 ml). The combined organic layers were washed with a saturated aqueous solution of NaHCO_3_ (100 ml). The organic layer was separated, dried over Na_2_SO_4_, filtered and concentrated under reduced pressure to afford 1-bromo-3-[1-(trifluoromethyl)cyclobutyl]benzene (4.0 g, 95.0% purity, 13.62 mmol, 69.2% yield).

^1^H NMR (400 MHz, CDCl_3_) δ 7.45 (dt, J = 7.2, 2.0 Hz, 1H), 7.43 (s, 1H), 7.30 – 7.20 (m, 2H), 2.76 (tdd, J = 10.4, 5.2, 2.4 Hz, 2H), 2.54 (q, J = 10.0 Hz, 2H), 2.22 (h, J = 9.0 Hz, 1H), 1.95 (tq, J = 10.3, 5.2 Hz, 1H).

### Step 2 of CBB1 synthesis

The Synthesis of 3-[1-(trifluoromethyl)cyclobutyl]benzoic acid (3): 1-Bromo-3-[1-(trifluoromethyl)cyclobutyl]benzene (3.5 g, 12.59 mmol) was dissolved in dry THF (100 ml). The reaction mixture was cooled to -78°C under Ar atmosphere. Then n-butyllithium (23% solution in hexane, 1.01 g, 1.48 ml, 15.74 mmol, 1.25 eq.) was added dropwise to the reaction solution at -78°C. Then the reaction mixture was stirred for 30 min at -78°C and poured onto an excess solid CO_2_. The mixture was allowed to warm to room temperature, stirred for 12 h and then concentrated in vacuo. Then the obtained residue was dissolved in water (100 ml) and washed with EtOAc (3 x 20 ml). The aqueous layer was separated, acidified to pH = 3 and extracted with EtOAc (3 x 75 ml). The combined organic extracts were dried over Na_2_SO_4_, filtered and concentrated under reduced pressure to afford 3-[1-(trifluoromethyl)cyclobutyl]benzoic acid (1.9 g, 95.0% purity, 7.39 mmol, 58.7% yield).

^1^H NMR (400 MHz, DMSO) δ 13.03 (s, 1H), 7.92 (d, J = 7.6 Hz, 1H), 7.84 (s, 1H), 7.55 (h, J = 6.1 Hz, 2H), 2.72 (ddd, J = 14.8, 9.6, 5.1 Hz, 2H), 2.58 (q, J = 9.9 Hz, 2H), 2.11 (q, J = 9.1 Hz, 1H), 2.01 – 1.85 (m, 1H).

### Step 3 of CBB1 synthesis

The Synthesis of 2-hydroxy-N-(4-methyl-2-oxo-2H-chromen-7-yl)-5-(3-(1-(trifluoromethyl)-cyclobutyl)benzamido)benzamide (CBB1): 3-[1-(Trifluoromethyl)cyclobutyl]benzoic acid (1.89 g, 7.73 mmol, 1 eq.) and 1-(1H-imidazole-1-carbonyl)-1H-imidazole (1.25 g, 7.73 mmol, 1 eq.) were dissolved in dry DMF (30 ml) and stirred at 75°C for 2 h. Then reaction mixture was cooled to ambient temperature and 5-amino-2-hydroxy-N-(4-methyl-2-oxo-2H-chromen-7-yl)benzamide (2.4 g, 7.73 mmol, 1 eq.) was added. The reaction mixture was stirred at ambient temperature for 12 h. Then the reaction mixture was diluted with water (50 ml), the resultant precipitate was filtered and washed with water (2 x 15 ml). The precipitate was dried under reduced pressure and subjected to FC (CHCl_3_/MeCN, 1:0 to 0:1) to give two fractions of 2-hydroxy-N-(4-methyl-2-oxo-2H-chromen-7-yl)-5-(3-(1-(trifluoromethyl)cyclobutyl)benzamido)benzamide: 1.1 g (100% purity, 2.05 mmol, 26.5% yield) and 0.85 g (86% purity, 1.36 mmol, 17.6% yield) as yellow solid.

_1_H NMR (400 MHz, DMSO) δ 11.25 (s, 1H), 10.81 (s, 1H), 10.30 (s, 1H), 8.17 (s, 1H), 7.98 – 7.88 (m, 3H), 7.83 (d, J = 9.1 Hz, 1H), 7.77 (d, J = 8.8 Hz, 1H), 7.65 (d, J = 8.4 Hz, 1H), 7.59 – 7.47 (m, 2H), 7.02 (d, J = 9.0 Hz, 1H), 6.30 (d, J = 1.5 Hz, 1H), 2.69 (d, J = 24.4 Hz, 4H), 2.42 (d, J = 1.3 Hz, 3H), 2.11 (s, 1H), 2.02 – 1.87 (m, 1H).

### CCB1 Alkylation

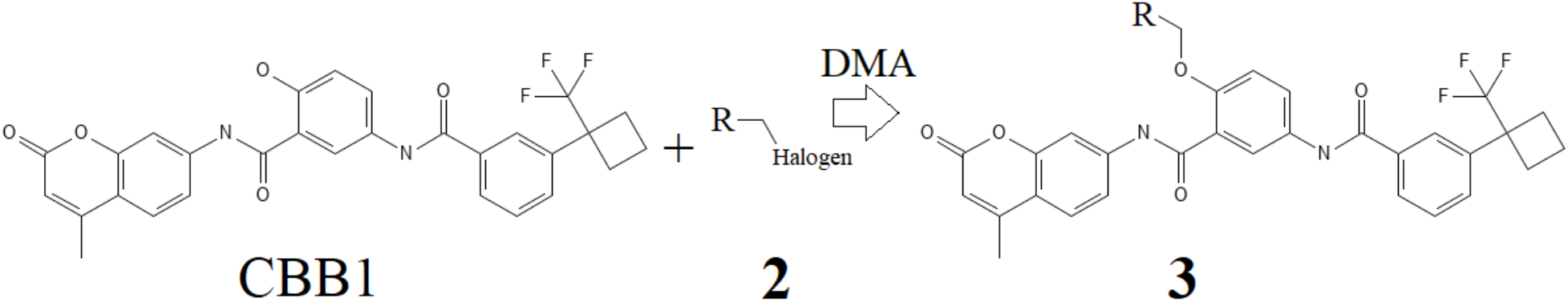

Reagent A – CBB1 – (1 eq.), Reagent B (1.3 eq.), Cs_2_CO_3_ (2.5 eq.) were mixed in dry DMA (appr. 0.7 ml per 100 mg of product). In case of using a salt of any of reagents, an additional amount of Cs_2_CO_3_ was added to the reaction mixture to transfer the reagent to the base form. The mixture was sealed and stirred at 70°C for 16 hour(-s). The solvent was evaporated under reduced pressure, and the residue was dissolved in DMSO (appr. 1 ml up to 100 mg of product). The solution was filtered, analyzed by LCMS, and transferred for HPLC purification.

The purification was performed using Agilent 1260 Infinity systems equipped with DAD and mass-detector. Waters Sunfire C18 OBD Prep Column, 100 A, 5 µm, 19 mm × 100 mm with SunFire C18 Prep Guard Cartridge, 100 A, 10 µm, 19 mm × 10 mm was used. Deionized Water (phase A) and HPLC-grade Methanol or Acetonitrile (phase B) were used as an eluent. In some cases, ammonia or TFA was used as an additive to improve the separation of the products. In these cases, free bases and TFA salts of the products were formed respectively.

### CCB1 Arylation

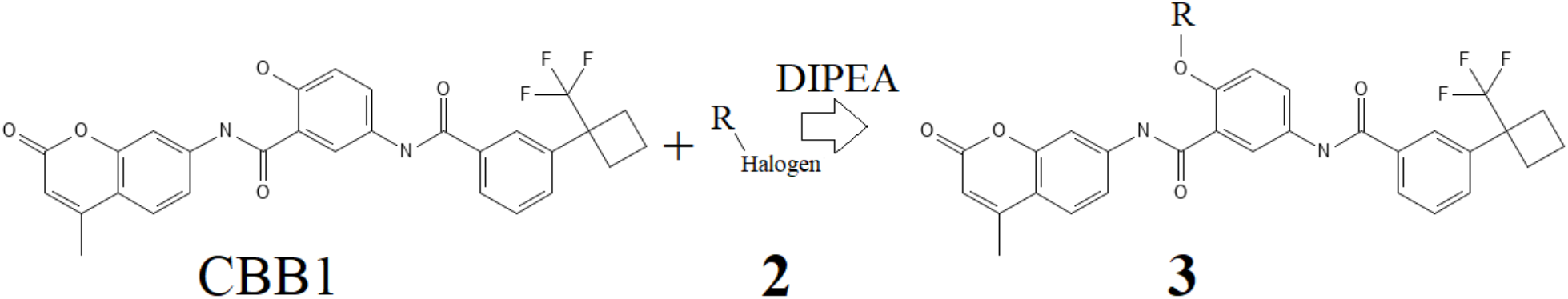

Reagent A – CBB1 – (1 eq.), Reagent B (1.2 eq.), DIPEA (3 eq.) were mixed in dry DMSO (appr. 2 ml per 100 mg of product). In case of using a salt of any of reagents, an additional amount of DIPEA was added to the reaction mixture to transfer the reagent to the base form. The mixture was sealed and stirred at 100°C for 18 hour(-s). After confirmation of completion of the reaction by LCMS the solution was filtered and transferred for HPLC purification.

The purification was performed using Agilent 1260 Infinity systems equipped with DAD and mass-detector. Waters Sunfire C18 OBD Prep Column, 100 A, 5 µm, 19 mm × 100 mm with SunFire C18 Prep Guard Cartridge, 100 A, 10 µm, 19 mm × 10 mm was used. Deionized Water (phase A) and HPLC-grade Methanol or Acetonitrile (phase B) were used as an eluent. In some cases, ammonia or TFA was used as an additive to improve the separation of the products. In these cases, free bases and TFA salts of the products were formed respectively.

### Synthesis of 2-hydroxy-N-(4-methyl-2-oxo-2H-chromen-7-yl)-5-[(4-phenylpiperidin-1-yl)-sulfonyl]-aminobenzamide (CBB2)

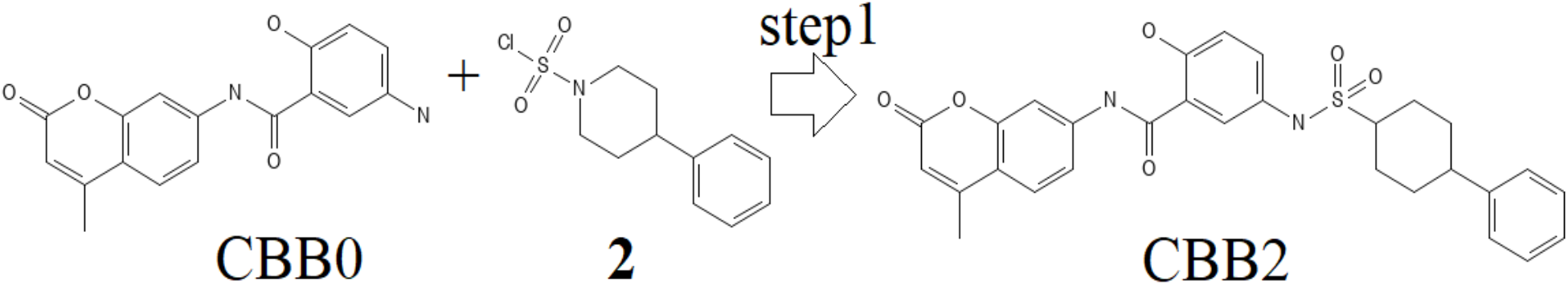

### Step 1 of CBB2 synthesys

Synthesis of 2-hydroxy-N-(4-methyl-2-oxo-2H-chromen-7-yl)-5-[(4-phenylpiperidin-1-yl)sulfonyl]aminobenzamide (CBB2). 5-Amino-2-hydroxy-N-(4-methyl-2-oxo-2H-chromen-7-yl)benzamide – CBB0 – (1.0 g, 3.22 mmol, 1 eq.) was suspended in dry pyridine (30 ml) and 4-phenylpiperidine-1-sulfonyl chloride (1.67 g, 6.45 mmol, 2 eq.) was added. The reaction mixture was stirred at ambient temperature for 72 h. Then the volatiles were evaporated under reduced pressure to give a crude product which was purified by FC (CHCl3/MeCN, 1:0 to 0:1) to give 2-hydroxy-N-(4-methyl-2-oxo-2H-chromen-7-yl)-5-[(4-phenylpiperidin-1-yl)sulfonyl]aminobenzamide (200.0 mg, 95.0% purity, 356.08 µmol, 11% yield) as a yellow solid.

1H NMR (500 MHz, DMSO) δ 11.23 (s, 1H), 10.73 (s, 1H), 9.66 (s, 1H), 7.92 (d, J = 2.1 Hz, 1H),7.78 – 7.73 (m, 2H), 7.63 (dd, J = 8.6, 2.1 Hz, 1H), 7.31 (dd, J = 8.8, 2.8 Hz, 1H), 7.22 (t, J = 7.6 Hz, 2H), 7.14 (dd, J = 7.9, 2.1 Hz, 3H), 6.99 (d, J = 8.8 Hz, 1H), 6.29 (d, J = 1.4 Hz, 1H), 3.69 (d, J = 12.2 Hz, 2H), 2.77 (dd, J = 13.3, 10.9 Hz, 2H), 2.56 (d, J = 12.4 Hz, 1H), 2.41 (d, J = 1.3 Hz, 3H), 1.73 (d, J = 12.5 Hz, 2H), 1.52 – 1.41 (m, 2H).

### CCB2 Arylation

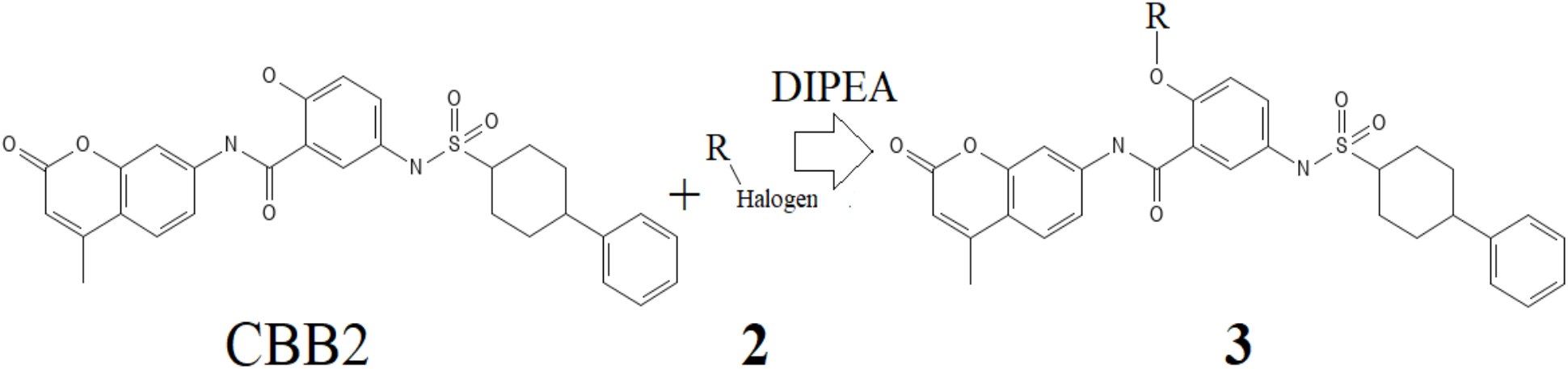

Reagent A – CBB2 – (1 eq.), Reagent B (1.2 eq.), DIPEA (3 eq.) were mixed in dry DMSO (appr. 2 ml per 100 mg of product). In case of using a salt of any of reagents, an additional amount of DIPEA was added to the reaction mixture to transfer the reagent to the base form. The mixture was sealed and stirred at 100°C for 12 hour(-s). After confirmation of completion of the reaction by LCMS the solution was filtered and transferred for HPLC purification.

The purification was performed using Agilent 1260 Infinity systems equipped with DAD and mass-detector. Waters Sunfire C18 OBD Prep Column, 100 A, 5 µm, 19 mm × 100 mm with SunFire C18 Prep Guard Cartridge, 100 A, 10 µm, 19 mm × 10 mm was used. Deionized Water (phase A) and HPLC-grade Methanol or Acetonitrile (phase B) were used as an eluent. In some cases, ammonia or TFA was used as an additive to improve the separation of the products. In these cases, free bases and TFA salts of the products were formed respectively.

#### LG-250

1H NMR (500 MHz, DMSO-d6) δ 13.00 – 12.53 (m, 1H), 11.44 (s, 1H), 11.21 (s, 1H), 10.82 (s, 1H), 8.26 (dd, J = 17.9, 2.3 Hz, 2H), 8.05 (ddd, J = 8.8, 6.6, 2.4 Hz, 2H), 7.99 (d, J = 8.6 Hz, 1H), 7.88 (d, J = 8.8 Hz, 2H), 7.70 (d, J = 8.9 Hz, 2H), 7.54 (d, J = 8.6 Hz, 1H), 7.25 (d, J = 8.9 Hz, 1H), 7.00 (d, J = 8.7 Hz, 2H), 6.26 (d, J = 1.5 Hz, 1H), 2.38 (s, 3H).

#### LG-273

1H NMR (500 MHz, DMSO-d6) δ 12.78 (s, 1H), 12.39 (s, 1H), 11.34 (s, 1H), 10.83 (s, 1H), 8.23 (d, J = 2.6 Hz, 1H), 8.05 (dd, J = 8.9, 2.7 Hz, 1H), 7.88 (d, J = 8.8 Hz, 2H), 7.71 (s, 1H), 7.70 (s, 1H), 7.55 (d, J = 8.7 Hz, 1H), 7.25 (d, J = 8.9 Hz, 1H), 7.00 (d, J = 8.7 Hz, 2H), 6.26 (d, J = 1.4 Hz, 1H), 4.32 (q, J = 7.1 Hz, 2H), 2.75 (s, 2H), 2.67 (s, 2H), 2.38 (s, 3H), 1.74 (d, J = 5.6 Hz, 4H), 1.33 (t, J = 7.1 Hz, 3H).

#### LG-306

1H NMR (500 MHz, DMSO) δ 12.78 (s, 1H), 11.33 (s, 1H), 10.83 (s, 1H), 10.06 (s, 1H), 8.45 (s, 1H), 8.23 (s, 1H), 8.04 (d, J = 8.9 Hz, 1H), 7.88 (d, J = 8.3 Hz, 2H), 7.78 (s, 1H), 7.70 (d, J = 8.2 Hz, 2H), 7.54 (d, J = 8.5 Hz, 1H), 7.25 (d, J = 8.9 10 Hz, 1H), 7.01 (d, J = 8.3 Hz, 2H), 6.26 (s, 1H), 3.98 (s, 3H), 2.38 (s, 3H).

#### LG-310

1H NMR (500 MHz,DMSO) δ 11.6 (s, 1H), 11.36 (s, 1H), 10.85 (s, 1H), 8.84 (d, J = 1.4 Hz, 1H), 8.48 (s, 1H), 8.26 (s, 1H), 8.12 - 8.01 (m, 1H), 7.9 (d, J = 6.9 Hz, 2H), 7.79 - 7.69 (m, 2H), 7.58 (s, 1H), 7.27 (d, J = 9.1 Hz, 1H), 7.03 (d, J = 7.1 Hz, 2H), 6.28 (s, 1H), 3.49 - 3.47 (m, 3H), 2.4 (s, 4H)

### *In vitro* Biological Assay

To assess the inhibitory effects of the synthesized compounds on the LAG-3/MHCII interaction, we employed the Homogeneous Time-Resolved Fluorescence (HTRF) LAG-3/MHCII binding assay (Revvity Inc., USA). This assay quantifies the interaction between biotinylated LAG-3 and MHCII using Streptavidin-Europium cryptate as the HTRF donor (binding LAG-3) and a d2-labeled antibody as the HTRF acceptor (binding MHCII). In the absence of an inhibitor, LAG-3 and MHCII interact, bringing the donor and acceptor fluorophores into close proximity. Upon excitation at 320 nm, the donor emits fluorescence at 620 nm, which is transferred via Förster Resonance Energy Transfer (FRET) to the acceptor, generating a secondary emission at 665 nm. The intensity of this 665 nm emission is directly proportional to the extent of LAG-3/MHCII complex formation. Thus, a higher signal indicates greater binding, whereas a reduced signal reflects inhibition of the interaction by test compounds.

For the assay, 2 μL of each test compound was dispensed into individual wells of an HTRF 96-well low-volume plate (PerkinElmer), followed by the sequential addition of 4 μL of LAG-3, 4 μL of MHCII, and 10 μL of a 1:1 (v/v) mixture of Streptavidin-Europium cryptate and d2-labeled antibody. The plate was incubated at room temperature for 3 hours before fluorescence measurements were recorded using a Tecan Spark plate reader. Donor and acceptor emissions were detected at 620 nm and 665 nm, respectively.

### Bioluminescent Cell-Based Assay

To assess the ability of compounds to inhibit the LAG-3/MHCII interaction in a cell-based system, we utilized the LAG-3/MHCII Blockade Bioassay (Promega Inc., USA) following manufacturer’s instructions. In brief, MHCII-positive cells were thawed in cell recovery medium (90% DMEM + 10% FBS) supplemented with the TCR-activating antigen and plated in two white 96-well assay plates at 100 µL per well. The plates were incubated at 37°C with 5% (vol/vol) CO_2_ for 20–24 hours. After incubation, the medium was removed, and serial dilutions of the control antibodies, Clone 17B4 and Relatlimab (Selleck Chemicals LLC., USA), or compounds, prepared in assay buffer (90% RPMI + 10% FBS), were added to the wells at 40 µL per well.

Jurkat T cells were then thawed in assay buffer, dispensed into wells at 40 µL per well, and incubated at 37°C with 5% (vol/vol) CO_2_ for 5 hours. Following incubation, 80 µL of Bio-Glo Reagent was added to each well, and luminescence was measured after a 10-minute incubation at room temperature using a Tecan Spark plate reader.

## Supporting information

Complex of CNMHTPMVC with LAG-3 after 1 us MD simulation

A refined LAG-3 (D1, D2) fragment after 10 us MD simulation

## Acknowledgements

The biological assays in this work were contractually performed by the Life Sciences Institute Biofactorial High Through-put Biology facility, which is supported by the UBC GREx Biological Resilience Initiative.

## Supplementary Information

- Additional pdb structures. lag3_D1D2_after_lmd.pdb - A refined LAG-3 (D1, D2) fragment after 10 us MD simulation lag3_CNMHTPMVC_after_lmd.pdb - Complex of CNMHTPMVC with LAG-3 after 1 us MD simulation

